# Opsonization by non-neutralizing antibodies can confer protection to SARS-CoV-2 despite Spike-dependent modulation of phagocytosis

**DOI:** 10.1101/2021.10.14.464464

**Authors:** Wael Bahnan, Sebastian Wrighton, Martin Sundwall, Anna Bläckberg, Urban Höglund, Olivia Larsson, Hamed Khakzad, Magdalena Godzwon, Maria Walle, Elizabeth Elder, Lotta Happonen, Oscar André, Johannes Kumra Ahnlide, Thomas Hellmark, Vidar Wendel-Hansen, Robert PA Wallin, Johan Malmström, Lars Malmström, Mats Ohlin, Magnus Rasmussen, Pontus Nordenfelt

**Affiliations:** Lund University, Faculty of Medicine, Department of Clinical Sciences Lund, Infection Medicine, SE-22184 Lund, Sweden; Infectious Disease Clinic, SUS, Lund, Sweden; Adlego Biomedical AB, Uppsala, Sweden; Equipe Signalisation Calcique et Infections Microbiennes, Ecole Normale Superieure Paris-Saclay, 91190 Gif-sur-Yvette, France; Institut National de la Sante et de la Recherche Medicale (INSERM) U1282, 91190 Gif-sur-Yvette, France; Lund University, Department of Immunotechnology, Lund, Sweden; Public Health Agency of Sweden, 17182 Solna, Sweden; Lund University, Skane University Hospital, Department of Clinical Sciences Lund, Nephrology, Lund, Sweden; Tanea Medical Ab, Uppsala, Sweden; SciEd Solutions, Stockholm, Sweden; Institute for Computational Science, University of Zurich, Winterthurerstrasse 190, CH-8057 Zurich, Switzerland

**Keywords:** SARS-CoV-2, Spike protein, antibody binding, antibody function, phagocytosis, in vivo model

## Abstract

Spike-specific antibodies are central to effective COVID19 immunity. Research efforts have focused on antibodies that neutralize the ACE2-Spike interaction but not on non-neutralizing antibodies. Antibody-dependent phagocytosis is an immune mechanism enhanced by opsonization, where typically, more bound antibodies trigger a stronger phagocyte response. Here, we show that Spike-specific antibodies, dependent on concentration, can either enhance or reduce Spike-bead phagocytosis by monocytes independently of the antibody neutralization potential. Surprisingly, we find that both convalescent patient plasma and patient-derived monoclonal antibodies lead to maximum opsonization already at low levels of bound antibodies and is reduced as antibody binding to Spike protein increases. Moreover, we show that this Spike-dependent modulation of opsonization seems to affect the outcome in an experimental SARS-CoV-2 infection model. These results suggest that the levels of anti-Spike antibodies could influence monocyte-mediated immune functions and propose that non-neutralizing antibodies could confer protection to SARS-CoV-2 infection by mediating phago-cytosis.

## Introduction

COVID19, caused by the SARS-CoV-2 virus, has since the end of 2019 resulted in millions of deaths and serious societal health effects. Treatment of patients with convalescence plasma or monoclonal antibodies was attempted early on during the pandemic, inspired by previous partial successes with Respiratory Syncytial Virus (1) and Ebola (2). Two monoclonal antibody cocktails targeting the SARS-CoV-2 Spike protein (casirivimab and imdevimab) (3) and (bamlanivimab and etesevimab) (4, 5) were given emergency use authorization by the FDA after positive phase III clinical trial data. Trials showed that antibody cocktails reduced symptoms, hospitalization, and mortality associated with COVID19 for early-stage infections. However, studies regarding their use for treating severe COVID19 showed no clinical benefit (6).

The therapeutic antibodies described previously neutralize the interaction between the Spike protein and the ACE2 receptor, thereby hindering viral entry into host cells. Considerable efforts have been made to generate neutralizing anti-Spike antibodies (7, 8, 9, 10). Neutralizing antibodies, however, constitute only a fraction of the antibody repertoire generated by B cells against the Spike protein during COVID19 infection (11). The opsonic capability has not been a focal point in the characterization of neutralizing antibodies. Non-neutralizing antibodies, comprising the majority of the humoral immune response to a pathogen, have other immunological functions such as complement-dependent immune activation and viral phagocytosis (reviewed by Forthal (12)). Phagocytosis plays a substantial role in the anti-viral immune response (13). Through virion or cellular phagocytosis, phagocytic cells help reduce the viral load by eliminating infection sources. In this context, we were interested in whether or not Spike antibodies might mediate phagocytosis as has been previously seen with influenza (13, 14, 15).

However, in other viral infections (such as Dengue, SARS-CoV-2, Respiratory Syncytial Virus, and others), insufficient levels of neutralizing antibodies allow non-neutralizing antibodies to mediate the entry of virions into host immune cells (16). This infection of immune cells via FcγR leads to Antibody-Dependent-Enhancement (ADE), exacerbating the infection and worsening patient outcomes (17). So far, studies on COVID19 vaccines and monoclonal antibodies utilized in COVID19 therapy have seen no evidence of ADE (16, 17, 18, 19, 20, 21). This clinical absence of ADE remains true even when some studies report that patient sera with high titers of neutralizing antibodies could induce Spike-bead phagocytosis or FcγR-activation (ADCP) (22, 23, 24).

Our work shows evidence that convalescent patient plasma and monoclonal anti-Spike antibodies induce phagocytosis but with diminishing returns when the antibody concentrations become high. We also demonstrate that the activation and inhibition of phagocytosis are independent of neutralization potential. Finally, we present data from an experimental animal infection model showing that non-neutralizing antibodies can protect animals from SARS-CoV-2 infection. The results in this study shed light on the importance of non-neutralizing antibodies in mediating phagocytosis and how their presence translates into protection after experimental infection.

## Results

### Convalescent patient plasma reduces Spike-mono-cyte interaction

Blood plasma was obtained from 20 COVID19 convalescent patients (Supp. Table 1). We used biotinylated Spike protein conjugated to streptavidin fluorescent microspheres (1 µm beads) as a model for Spike-monocyte interactions. The beads were used as bait for THP-1 monocytes. To opsonize the beads, we incubated them with the patient plasma at different dilution levels. We chose the 0.01-1% concentrations to mimic IgG levels in the mucosal niche or tissues, which would be the first place of encounter with the SARS-CoV-2 virus. The highest level of association between plasma-opsonized Spike-beads and cells was at the intermediate plasma dilution (0.1%), while the higher and lower concentrations of plasma (1 and 0.01%, respectively) showed reduced association (Fig. 1a). In fact, the only consistent effect we saw across our patient plasma samples was a reduction in Spike-particle association with THP-1 cells at the highest plasma concentration. This phenomenon was seen in 18 out of 20 patient samples. Two patient samples (patients 8 and 18) showed no or low opsonic ability. The reduction in Spike-THP-1 cell association under high plasma concentrations was independent of patient sex, age, or disease severity (Supp. Table 1).

**Fig. 1.**
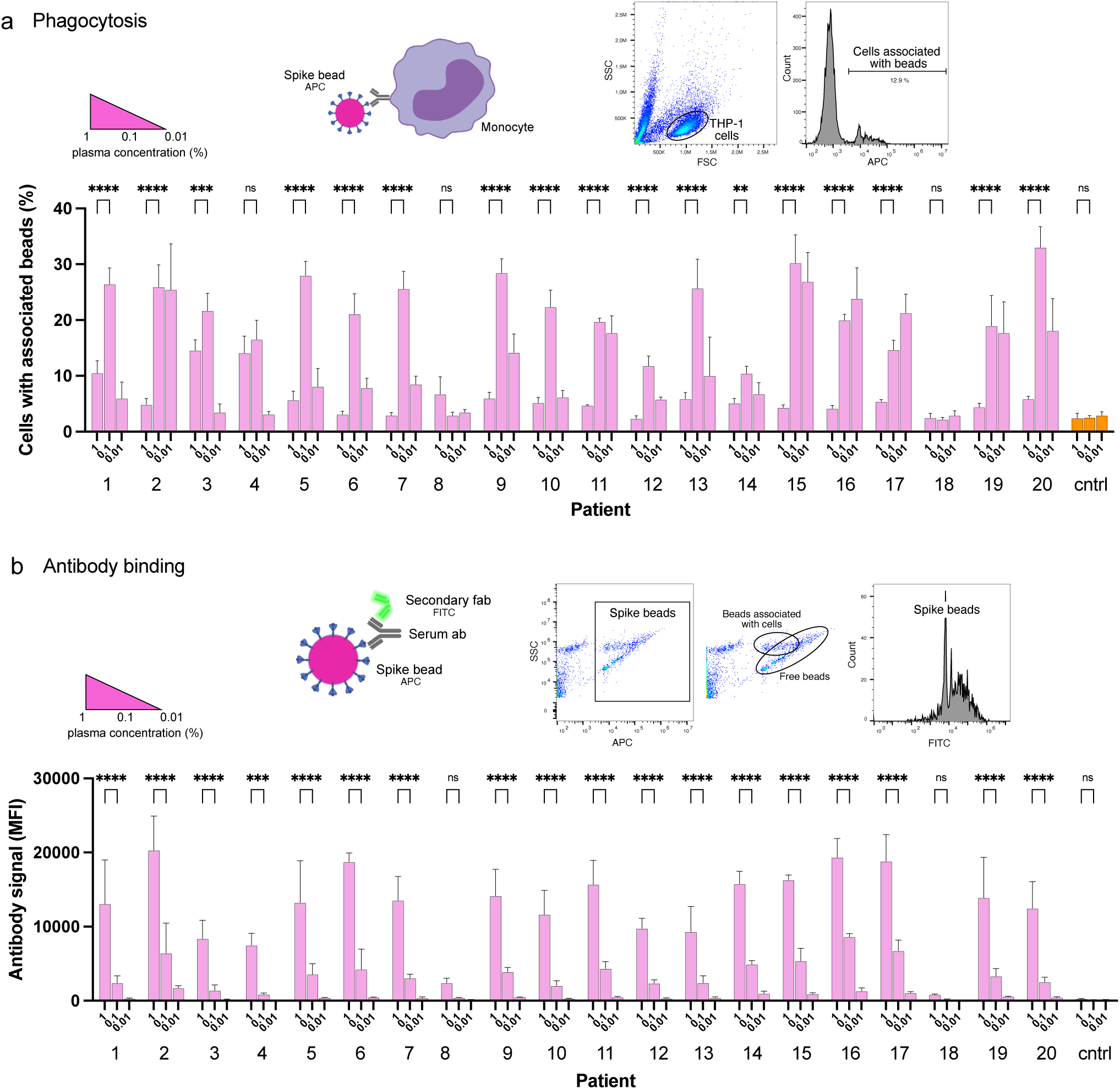
Convalescent patient plasma reduces Spike-monocyte interaction. **a** Biotinylated Spike protein was conjugated to fluorescent (APC) streptavidin microspheres and was opsonized with three convalescent patient plasma concentrations (1%, 0.1%, and 0.01%). The beads were then mixed with THP-1 cells at a ratio of 2:1, and the association was measured using flow cytometry. Cells that had signal in the APC channel were considered positive. The gating strategy is shown in the top right. **b** The same samples of THP-1 cells and beads from (a) were fixed with methanol and stained with a fluorescent (FITC) Fab anti-human Fab secondary antibody. The samples were analyzed for human antibody (opsonin) binding to the Spike-beads using flow cytometry. The gating strategy is shown in the top right. The data presented are from three independent experiments. Error bars represent the SD. Statistical significance was assessed using two-way ANOVA with Dunnett’s multiple comparison correction. * denotes p 0.05, ** for p 0.01, *** for p 0.001 and **** for p 0.0001).

As our results were unexpected, we checked whether the reduction in particle-to-cell association seen at higher plasma concentrations (1%) was due to a loss of Spike or antibody binding. For that purpose, we methanol-fixed the samples from the experiment shown previously (Fig. 1a). The samples were then stained with a fluorescently conjugated (FITC) secondary antibody (Fab anti-human Fab), which would react with the plasma anti-Spike antibodies which had bound to Spike on the beads. Unsurprisingly, increased plasma concentrations led to increased binding of Spike-specific antibodies to the Spike-beads (Fig. 1b). In contrast, patients 8 and 18 showed no or very low binding of antibodies to Spike-beads, correlating with overall reduced opsonization (Fig. 1a). Our results show that when assayed at higher concentrations, patient plasma is not permissive to THP-1 cell-Spike interactions, despite having antibodies that readily bind Spike protein.

### Generation of Spike-reactive human monoclonal antibodies

Considering our previous data showing that high concentrations of COVID19 convalescent plasma reduced Spike-THP-1 cell interactions compared to low concentrations, we decided to identify the role monoclonal antibodies play in Spike-THP-1 cell interactions. We isolated Spike-reactive B cells from convalescent COVID19 patients and performed single-cell sequencing (Fig. 2a). We chose 96 antibodies for production that were equidistantly spaced on the genetic clustering tree (Supp. Fig. 1a). The antibodies were expressed in HEK293 cells. ELISA-based screening of the antibody-containing supernatants allowed us to identify ten Spike-reactive antibodies (Fig. 2b, Supp. Fig. 1b-c), which belonged to different IgG germlines (Supp. Fig. 1d). The Spike-reactive antibodies were then assayed for reactivity against Spike-beads using flow cytometry, where we observed that nine antibodies were reactive to the Spike-beads (Fig. 2c). Ab11, 57, 59, 66, 77, 81, 94, and 95 showed clear reactivity (>40% positive beads) when assayed with Spike-beads at a concentration of 1 µg/ml. Ab59 demonstrated strongest binding, as could be seen through the relative increase in bead staining. Xolair (used at 10 µg/ml) and normal (pre-COVID19) plasma served as negative controls, whereas COVID19 plasma from a convalescent patient was our positive control.

**Fig. 2.**
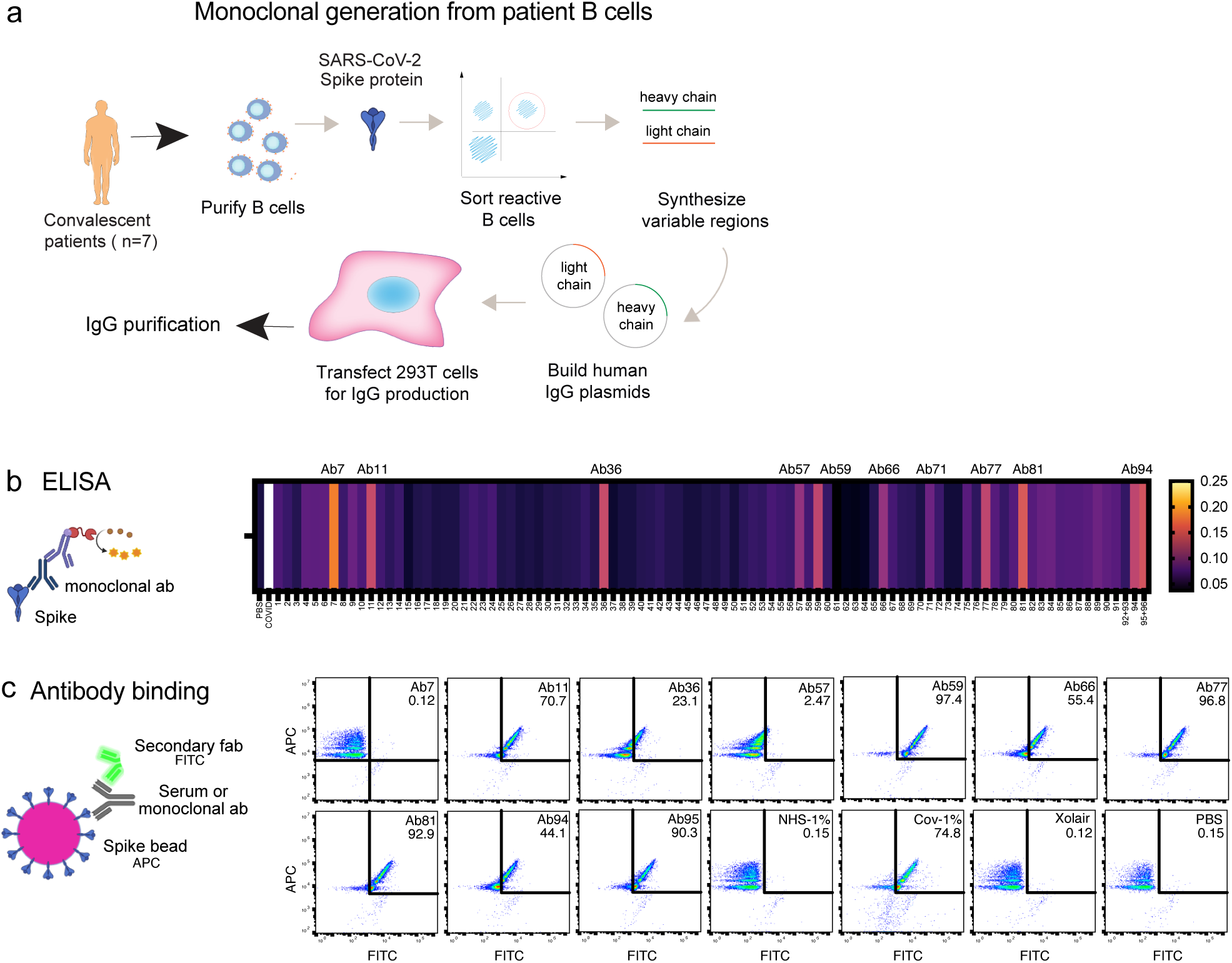
Generation of Spike-reactive human monoclonal antibodies. **a** Human monoclonal antibodies were generated from convalescent donor B cells through single-cell sequencing technology. 96 antibodies derived from Spike-reactive human B cells were produced in HEK293F cells. **b** Cell culture supernatants containing the antibodies were assayed by ELISA for reactivity against immobilized Spike protein. Serum from a COVID19 patient was used as a positive control. The data represent three replicate ELISAs where reproducibly reactive antibodies are indicated with their names above the heatmap. **c** Antibodies which were Spike-reactive in (b) were assayed for reactivity to Spike immobilized on beads. Fluorescent (APC) Streptavidin beads coated with biotinylated Spike protein were incubated with HEK293F-produced antibodies at a concentration of 1 µg/ml. The beads were then stained with a fluorescent (FITC) secondary anti-Fab antibody. The beads were analyzed by flow cytometry. Antibodies that shifted the beads into the FITC-positive gate were deemed reactive.

### Epitope mapping and structural mass spectrometry identify antibody binding sites

To identify antibody binding sites, we first used ELISA to study Spike domain interactions with RBD, RBD with L452R and T478K mutations (delta), and NTD from Spike (Fig. 3a). We could detect binding to seven antibodies, with high integrated signal (0.2-30 nM titration curves) for Ab59, Ab66, Ab81, and Ab94. Ab66 showed stronger interaction with delta RBD, and Ab 81 showed a lower signal. Ab94 only bound the NTD of Spike. We also performed relative antibody epitope mapping using the single-chain antibody fragments (scFv) isolated from an extensive combinatorial library. scFv mapping revealed that the Ab59 epitope overlaps with those of two scFv (A03_D03 and E01_C09, Fig. 3b) that interfere in the binding of Spike to ACE2.

**Fig. 3.**
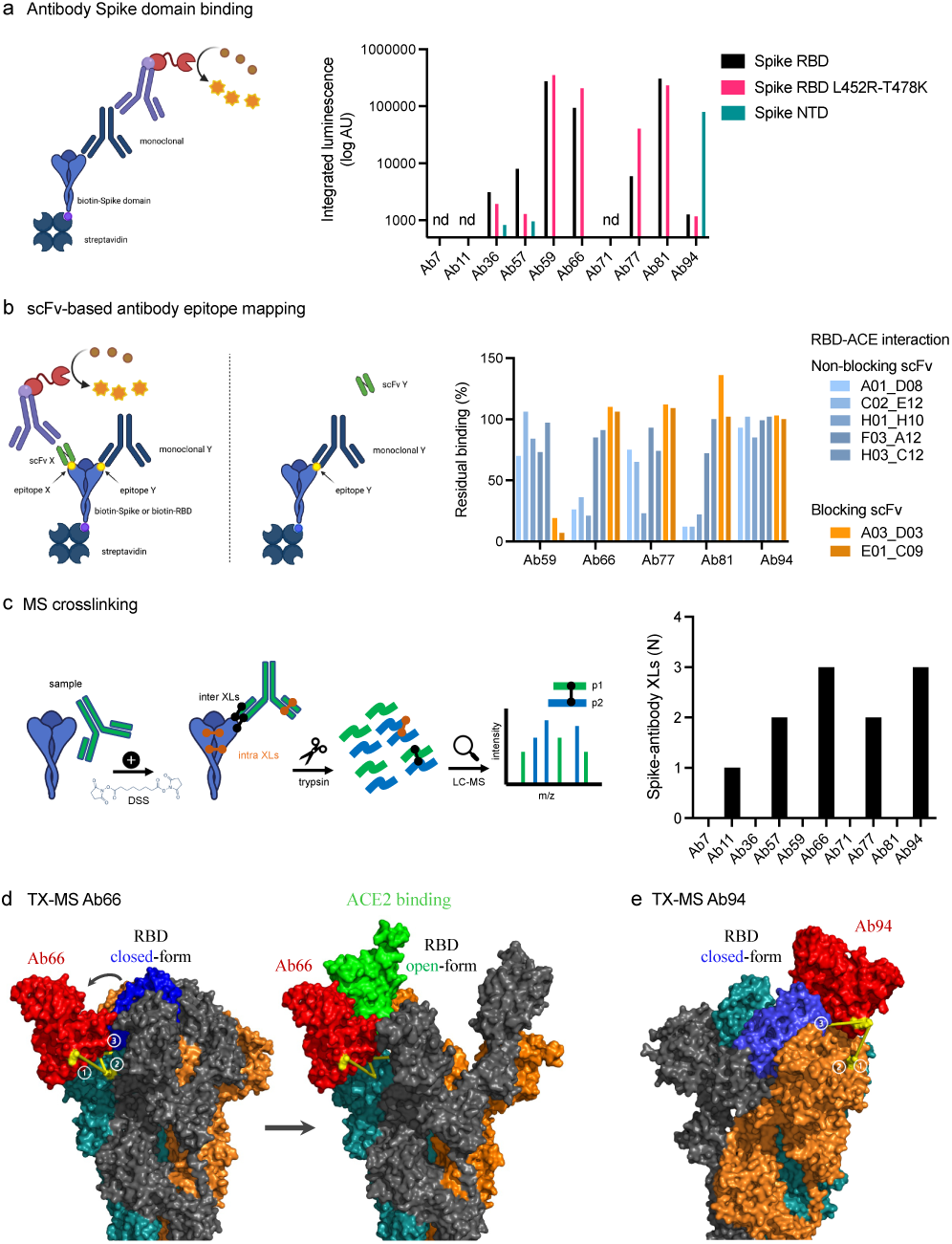
Epitope mapping and structural mass spectrometry identify antibody binding sites. **a** Antibody binding to Spike domains was analyzed using ELISA as shown with HRP signal as readout. Antibodies were titrated at 0.2-30 nM, and the integrated signal was calculated. The relative binding to each Spike domain is shown. nd = not determined. **b** Epitope mapping was performed using scFvs targeting RBD epitopes as shown. Antibody blocking of scFv binding was measured using anti-FLAG HRP signal. Representative of two independent experiments. **c** Spike protein was mixed with anti-spike antibodies and the complex was cross-linked with DSS, allowing for inter and intra cross-links. After trypsinization, mass spectrometric analysis was performed. The table to the right displays the number of inter-protein cross-links detected between Spike and its corresponding antibody. **d-e** The binding sites for Ab66 and Ab94 were determined by TX-MS using the cross-links from c, and the data was modelled using Rosetta. Models for Ab66 (d) binding the Spike protein in both its open and closed conformations as well Ab94 (e) are shown.

Next, we used TX-MS (25) to determine the binding interface between the SARS-CoV-2 specific antibodies and the RBD domain of the Spike protein. In short, we cross-linked the ten antibodies separately to the RBD domain, followed by mass spectrometry analysis and structural modeling (26). This resulted in the identification of 11 confident inter-protein XLs between the RBD domain and five of the antibodies (Ab11, Ab57, Ab66, Ab77, and Ab94) in addition to 30 intra RBD XLs (Fig. 3c, Supp Fig 3). The results show that the five antibodies can bind to the Spike protein, but they do not appear to compete with the binding site of human ACE2 directly. The interaction between Ab66 and Spike protein show binding to the open-state but not the closed-state (Fig. 3d). Further, the structural model indicates no competition between Ab66 and human ACE2, which is in accordance with previously published work, as only the open-state is responsible for binding human ACE225. In contrast, Ab94 appears to preferably bind the closed state (Fig. 3e). The combined data from our epitope analysis approaches indicate that Ab11, 57, 59, 66, 77, 81 bind Spike RBD, that Ab94 could interact with both RBD and NTD, and that Ab59 could be a neutralizing antibody, whereas the others are likely non-neutralizing.

### Neutralization assays identify one monoclonal which blocks the ACE2-Spike protein interaction

Typically, the most important biological function attributed to antibodies in the context of a viral infection is neutralization. We assayed our Spike-reactive antibodies for Spike-neutralization using three different approaches: Spike RBD-ACE2 protein binding (Fig. 4a), Spike particle-ACE2 cell interaction (Fig. 4b), and SARS-CoV-2 pseudovirus infection neutralization (Fig. 4c). The SPR-based Spike RBD-ACE2 binding data showed that Ab36 and Ab94 did not appear to interfere with RBD-ACE2 binding and that Ab57 reduced binding slightly. Ab59 completely blocked RBD-ACE binding, whereas Ab66, Ab77, and Ab81 seemed to bind well without interfering with the interaction. Next, we utilized ACE2-expressing HEK293 cells as a surrogate for lung epithelial cells. We measured the ability of Spike-beads to bind HEK293-ACE2+ cells after being opsonized with antibody supernatants. We assayed all 96 of our antibody-containing supernatants for Spike-particle neutralization. Only Ab59 showed a robust and reproducible reduction in Spike-particle binding to HEK293-ACE2+ cells compared to COVID19 patient plasma (Fig. 4b). As expected, pre-COVID19 plasma showed no inhibition of Spike-ACE2 interactions. Representative images of our experiments which were used for analysis, are shown in Supplementary Fig 2. We utilized a pseudovirus binding assay to verify the ability of our antibodies to neutralize SARS-CoV-2. Consistent with our previous experiments with RBD and Spike-beads (Fig 4a-b), we saw that Ab59 was the best (EC_50_: 19 ng/ml) among our antibodies in neutralizing SARS-CoV-2 pseudovirus infection (Fig. 4c). Taken together, our results indicate that of our 96 antibodies, only Ab59 is a potent neutralizer of the Spike-ACE2 interaction.

**Fig. 4.**
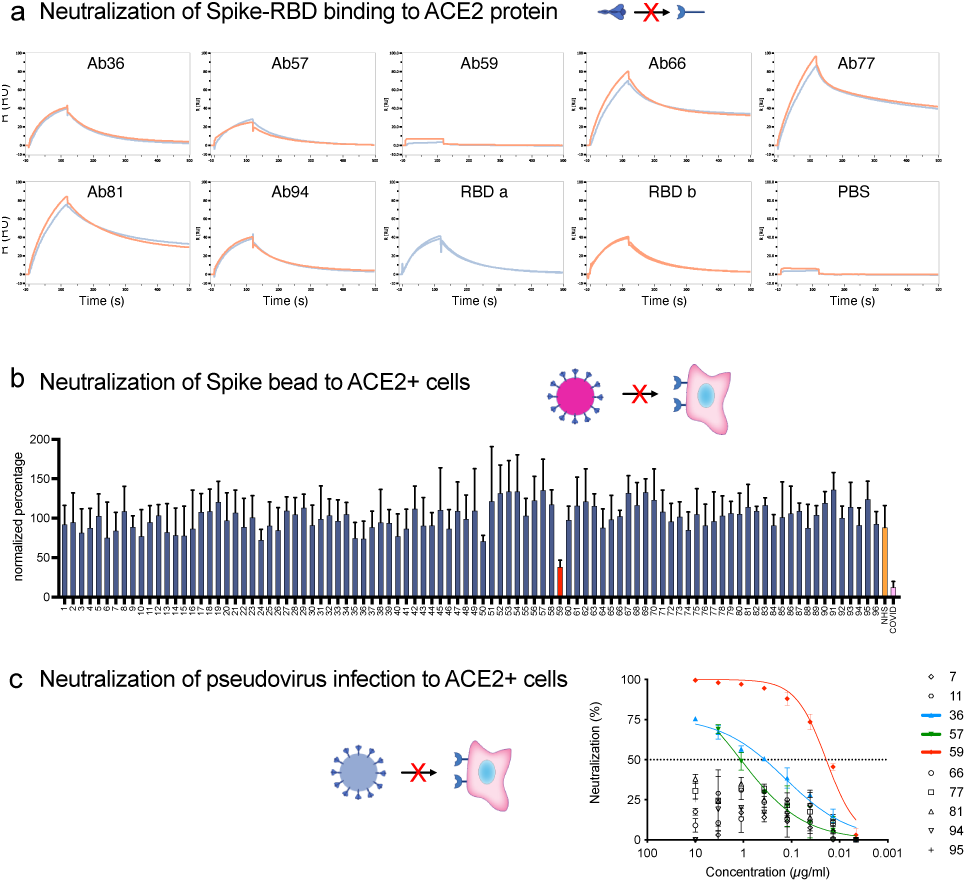
Neutralization assays identify one monoclonal as blocking the ACE2-Spike protein interaction. **a** SPR analysis of the binding of monoclonal antibodies to the RBD domain of the Spike protein. PBS served as a negative control, and the intact RBD was our positive control for ACE2 binding. **b** The 96 antibodies which we produced were assayed for neutralization potential in a Spike-bead-based neutralization assay. Spike-beads (such as the ones used in (b)) were opsonized with the antibodies in 96 well plates. The beads were then centrifuged, reconstituted in fresh media, and added to HEK293-ACE2 cells at a ratio of 20 beads per cell and imaged with automated microscopy. The data is from 4 pooled experiments and is presented as bead association normalized percentage. Error bars indicate the SEM for the replicate experiments. **c** The 10 Spike-ELISA reactive antibodies were assayed for pseudovirus neutralization. A firefly luciferase encoding pseudotype lentivirus was used to infect HEK239-ACE2 cells. Antibody serial dilutions were used to block the viral entry into the HEK293-ACE2 cells. Nonlinear regression lines were fitted for the three antibodies that showed a higher than 50% reduction of infectivity. Those antibodies were highlighted in green (Ab57), blue (Ab36), and red (Ab59).

### High levels of human monoclonal antibodies reduce Spike-monocyte interaction

Antibodies are the primary mediators of FcγR-dependent cellular interactions. Given our previous data that high concentrations of convalescent patient plasma can reduce Spike-bead association with THP-1 monocytes (Fig 1.), we tested whether this reduction was antibody-driven. We chose antibody concentrations that were in a similar range (100 - 0.01 µg/ml) than what is expected at the plasma concentrations used (1% - 0.1%) (Fig. 1) and included a higher plasma concentration for comparison (10%). Interestingly, as with patient blood plasma, serially diluted Spike-specific monoclonal antibodies showed the same inhibition trend of bead-to-cell association at the higher concentrations (Fig. 5a). This association was confirmed to reflect the internalization of particles (i.e., phagocytosis) by using a pH-dependent fluorescent dye (Supp. Fig. 4). Also, as with plasma, this inhibition was correlated with increased antibody binding to Spike (Fig. 5b). It is important to note here that among the antibodies, Ab94 seemed to have almost half the binding efficiency of Ab59, an attribute that will be central for other experiments. The neutralizing antibody, Ab59, showed the same trend as the other non-neutralizing antibodies. We have thus identified that Spike-specific monoclonal antibodies isolated from COVID19 patients modulate the Spike-THP-1 cell interactions in a dose-dependent manner. This phenomenon is independent of the Spike-ACE2 neutralization capability of the monoclonal antibodies.

**Fig. 5.**
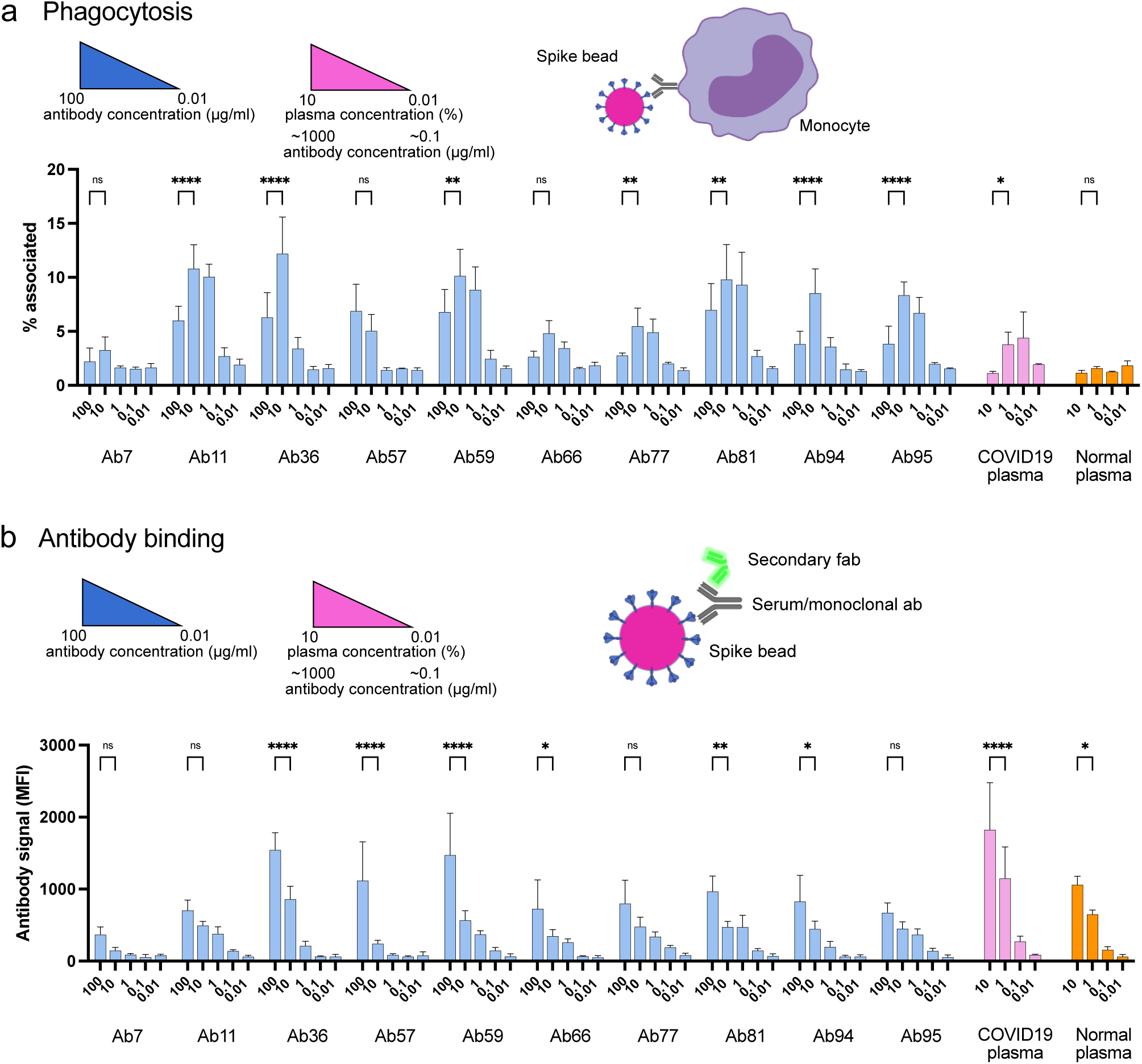
High levels of human monoclonal antibodies reduce Spike-monocyte interaction. **a** Spike-reactive monoclonal antibodies at concentrations of 100, 10, 1, 0.1, and 0.01 µg/ml were used to opsonize Spike-beads. Plasma was used at serial dilutions of 10%. The beads were then incubated with THP-1 cells at a ratio of 2 beads/cell. The cells were then analyzed by flow cytometry for association with the fluorescent Spike-beads. The data show the % of bead-associated cells and is pooled from three independent experiments. Error bars represent the SD. **b** The cells used in (a) were fixed with methanol and restained with a fluorescent (FITC) Fab anti-human secondary antibody. The samples were assessed for human antibody (opsonin) binding to the Spike-beads using flow cytometry. The data are from three independent experiments. Error bars represent the SD.

### Non-neutralizing antibodies can protect against SARS-CoV-2 infection

We have shown that among our antibodies, Ab59 is neutralizing while the other monoclonals are not. We have also demonstrated that our Spike-bead reactive antibodies are efficient at mediating Spike-mediated phagocytosis but reach a threshold after which there is a reduction in interaction efficacy. To test the antibodies’ function in a physiologically relevant context, we assessed different doses of neutralizing and non-neutralizing antibodies in an experimental animal infection model (Fig. 6a). We infected humanized ACE2 mice intranasally with 105 PFU (SARS-CoV-2; Wuhan strain from Swedish isolate). As a treatment model, we administered our monoclonal antibodies intraperitoneally a day after infection. Based on previous experience, we used the pseudovirus neutralization data (Fig. 4c) to calculate a protective dose in a prophylactic model (100 µg for Ab59). To test the effects of high dose administration, Ab59 was given at five times the calculated protective dose. For Ab94, we chose the same dose that would be considered protective for Ab59 (100 µg), as well as a higher dose (250 µg), which would be equivalent to the protective Ab59 dose based on the lower affinity of Ab94 (2.5 times lower, Fig. 2C and Fig. 5B). Interestingly, the best-protected animal group (lower weight loss) was the one where the animals were treated with the equivalent to a protective dose of our non-neutralizing yet opsonic Ab94 (Fig. 6b). Unexpectedly, the animals treated with a low dose of Ab59 fared better than the ones with the high dose, which had the worst outcome (more pronounced weight loss) among the treated groups. The low dose of Ab94 offered negligible improvement compared to untreated animals (Fig. 6b). The animal data indicate that too high doses of neutralizing antibodies are not beneficial in a treatment model and that non-neutralizing antibodies can offer protection to SARS-CoV-2 infection.

**Fig. 6.**
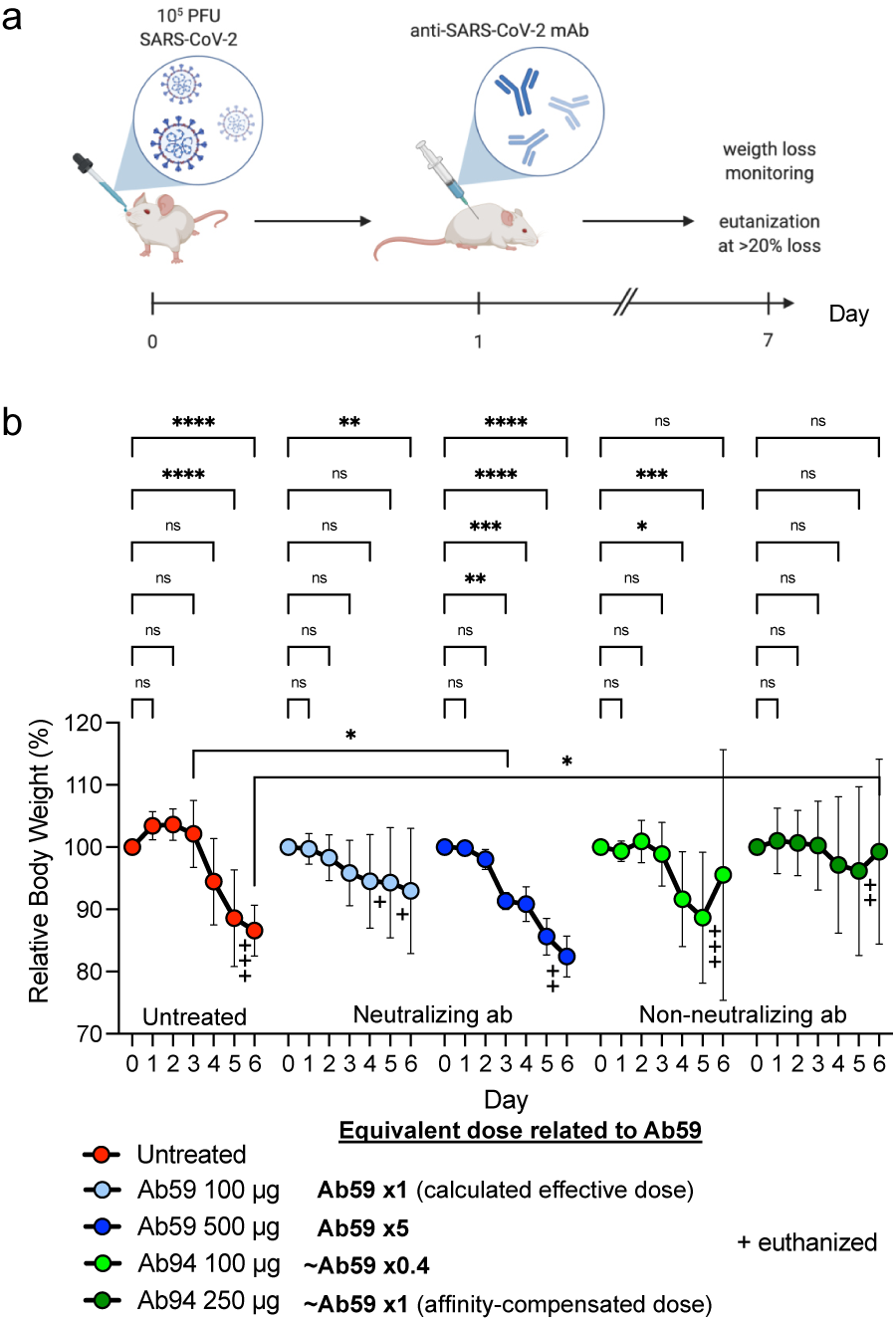
Non-neutralizing antibodies can protect against SARS-CoV-2 infection. **a** Humanized ACE2 mice were infected intranasally with SARS-CoV-2 (Wuhan strain). One day after infection, the animals (N=7 per group) were treated intraperitoneally with antibodies. Relative body weights were recorded and tabulated. **b** Body weights relative to each individual mouse over the time course of viral infection and treatment. Error bars represent the SD. Statistical significance was assessed using two-way ANOVA with Dunnett’s multiple comparison correction both within each treatment group and across the groups for each day. * denotes p 0.05, ** for p 0.01, *** for p 0.001 and **** for p 0.0001).

## Discussion

In this report, we present data on antibody modulation of Spike-monocyte interactions. To the best of our knowledge, only one previously published report showed Spike-bead phagocytosis after opsonization with 50% heat-inactivated serum and a 16 hours incubation of beads with THP-1 cells (22). We believe that the data from our experiments are more representative of the first few events after Spike-monocyte contact. Phagocytosis of small particles such as virions is a process that takes minutes, not hours (27). That is why we use shorter incubation times (30 min) and perform dose-response analysis across varying plasma concentrations. The dose-response analysis we performed exposed an antibody-mediated modulation of the Spike-monocyte interactions. It is important to elaborate on the concentrations we utilized, be it for plasma or monoclonal antibodies. For plasma, as tissues have a lower concentration of plasma proteins than whole blood, we used 1% as the highest concentration. As for the monoclonal antibody concentrations, we used 100 µg/ml as the highest concentration because it is roughly 1% of the antibody concentration present in plasma (10 mg/ml). Even though most of the experiments in this study are performed in vitro, we believe the effects we observe on phagocytosis could also be relevant in vivo, as the modulation effects occur already at relatively low antibody concentrations and would thus cover many physiological niches and scenarios.

That an increased binding of antibodies to a prey results in reduced or blocked phagocytosis is in stark contrast to what is typically expected (28). It cannot be explained by specific monoclonal interactions, as it is seen across diverse monoclonals as well as in convalescent polyclonal samples. This block in phagocytosis is only related to Spike protein. A combination of known mechanisms could potentially explain how SARS-CoV-2 could avoid phagocytosis using Spike protein. Bivalent trans-binding of antibodies is known to promote virion phagocytosis (29), where antigens are cross-linked depending on their density at the surface. Spike protein density on SARS-CoV-2 varies (30), and is increased with the D614G mutation (31). An increase in Spike antibody levels would lead to a competition of epitope binding, ultimately favoring the switch from bivalent trans-binding to monovalent binding, potentially leading to a reduction in phagocytosis. A synergistic mechanism could further aid SARS-CoV-2. Antigen height (especially below 10 nm) is important for efficient phagocytosis (32), and most likely, a consistent antigen height is beneficial as well. Besides altering its density, SARS-CoV-2 also appears to be able to dramatically change the Spike protein conformation, where some proteins stand up vertically from the surface, and others are tilted down horizontally (30). At high anti-Spike levels, this would present an approaching phagocyte with a monovalently opsonized, irregular surface with variable antigen height (15-25 nm), making the interaction difficult. In contrast, at low anti-Spike levels, the antibodies would be able to clasp Spike proteins in a bivalent, upright manner, presenting the phagocyte with a coherently opsonized surface at an effectively low antigen height. Careful mechanistic studies are needed to test this hypothesis.

How does the concentration-dependent phagocytosis modulation translate into an infection model? We utilized our anti-Spike antibodies as a therapeutic regimen in our animal infection. This creates a scenario where the mouse is already fighting the infection before therapy gets administered. The requirements, therefore, on therapeutic antibodies are higher than on prophylactic ones. When comparing the 100-µg dose of Ab59 and Ab94 (equivalent to 2.5 times lower binding affinity), we noticed that Ab59 showed better protection than the latter antibody. This could be attributed to either the neutralizing activity of Ab59, which Ab94 lacks, or the lower effective dose, given the lower Spike binding affinity of Ab94. Consistent with our data on phagocytosis (Fig. 5), animals treated with the high dose of Ab59 (500 µg) fared worse than the animals that got the 100-µg dose. Interestingly, the animals that got the affinity-corrected dose for Ab94 (250 µg, equivalent to 100 µg of Ab59 in terms of binding affinity) fared the best in our cohort. The animal infection data on Ab94 protecting the mice is, most importantly, congruent with the phagocytosis data, indicating a role in infection management for non-neutralizing antibodies. In future studies, it remains to be established how infections would proceed in animals treated with an excessive dose of non-neutralizing antibodies. Our animal experiments show that monoclonal antibodies could be viable therapeutics even if they lack neutralizing potential. This is an interesting discovery that, to the best of our knowledge, has not been shown before. Additionally, the dose variation data from Ab59-treated groups reflects the data we have presented in the concentration-dependent modulation of Spike phagocytosis section. These results may explain the clinical findings seen in antibody therapy trials in the sense that having low or excessive dosage of antibodies offers no clinical benefit (33).

It is widely accepted that a strong positive correlation exists between COVID19 disease severity and antibody titers (34, 35, 36, 37). High antibody titers are generally associated with severe disease and hospitalization. The high titers are thought to be a consequence of the severe infection. Our results pose a new question: could the high anti-Spike titers seen in hospitalized patients instead (at least partially) contribute to the immune dysregulation and worsening patient outcomes? These questions are relevant in the light of the FDA’s recommendation not to use monoclonal antibodies in hospitalized COVID19 patients (and who are seropositive) due to possible worsening of symptoms (3, 5). At the same time, it is important to note that convalescent plasma treatment for COVID19 was shown to be neither beneficial nor detrimental (38). Overall, the results presented in this study highlight a concentration-dependent modulation of phagocytosis by anti-Spike antibodies. This modulation phenomenon might help explain the unclear clinical benefit seen with monoclonal antibody treatment for COVID19. This modulation is seen in patient material and translates well to animal infection experiments. This opens several avenues for further experimentation. The biophysical mechanism underlying the antibody-mediated phagocytic modulation is an exciting topic to pursue, as are the bridging immune steps between phagocytosis and animal protection.

## Methods

### Cell culture, transfection, and protein production

THP-1 cells were maintained in RPMI medium supplemented with 10mM L-Glutamine and 10% fetal bovine serum (FBS). The cells were split kept at a density between 5×10^5^ and 10^6^ cells/ml. The cells were split when they reached a density of 10^6^ cells/ml, down to 5×10^5^ cells/ml. HEK293 cells (Sigma Aldrich 12022001-1VL) were maintained in DMEM supplemented with L-glutamine and 10% FBS. The cells were kept at 90% confluence and were not allowed to grow past passage 20. The plasmids for the 96 antibodies were aliquoted into 96 well plates. The cells to be transfected were grown in 24-well plates, with 500 µl of tissue culture medium. The plasmids were transfected into adherent HEK293 cells using the PEI method (39). The day after transfection the cell culture supernatant media was replaced with serum-free OptiMEM medium for 2 extra days. The supernatants containing the antibodies were distributed also in 96 well plates and stored for maximum one week for experimental use. HEK293 cells constitutively expressing the ACE2 receptor were acquired from BEI resources (NR-52511). They were maintained in DMEM supplemented with L-glutamine and 10% FBS for a maximum of 12 passages before being discarded. Expi293F suspension cells were purchased from Gibco (ThermoFisher) and routinely cultured in 125 ml Erlenmeyer flasks (Nalgene) in 30 ml Expi293 medium (Gibco) in an Eppendorf s41i shaker incubator at 37°C, 8% CO2, 120 rpm. Cells were passaged and split to a density of 0.5 × 106 cells/ml every 3 to 4 days. The day before transfection, the cells were seeded at a density of 2 × 10^6^ cells/ml. The next day, cells were seeded at 7.5 × 10^7^ cells in 25.5 ml Expi293 medium. The transient transfection was carried out using 100 µl of Expifectamine (Gibco) according to the manufacturer’s instructions. For Spike protein production, we used 40 µg of the Spike CS/PP plasmid (generously donated by Dr. Florian Krammer’s lab). For antibody production, 20 µg of plasmids for the heavy and light chain was used, respectively. For all plasmids, 16 hours after transfection, 150 µl of enhancer 1 and 1.5 ml of enhancer 2 (Expifectamine transfection kit, Gibco) were added and cells cultured for an additional 3 days. The cells were then pelleted at 400 x g, 5 min, RT and the supernatant transferred to new tubes. Magne Protein G beads (Promega) were used to purify the antibodies according to the manufacturer’s instructions.

### Antibody phage selections

SARS-CoV-2 Spike RBD-specific scFv were selected by phage display technology from a human synthetic scFv library(40), similar in design and construction to previously reported(41). Briefly, selection of specific binders was performed through a process similar to the one described in the past(41) using biotinylated proteins SARS-CoV-2 S1 protein, His, Avitag™ (ACRO Biosystems, S1N-C82E8) and SARS-CoV-2 Spike RBD, His, Avitag™ (ACRO Biosystems, SPD-C82E9) immobilized on paramagnetic beads (Dynabeads M-280 streptavidin; Invitrogen Dynal AS, Oslo, Norway) as target antigen. Phagemid DNA from the third and fourth rounds of phage selection was isolated (Plasmid Miniprep kit, Qiagen) and the genes encoding scFv fragments were ligated into an in-house constructed screening vector providing the secreted scFv with a triple-FLAG tag and a hexahistidine (His6) tag at the C-terminus. The constructs were subsequently transformed into TOP10 E. coli and individual, soluble scFv were produced as described elsewhere (41). Binding of individual selected scFv was initially assessed by ELISA against biotinylated antigen. Seven scFvs specific for RBD isolated this way were used to map relative epitope location of human IgG. The scFvs bind four epitopes on RBD, and two of them (A03-D03 and E01-C09) also interfere with the RBD-ACE2 interaction (data not shown).

### COVID19 patient samples and B cell isolation

For the spike-THP-1 association experiments, 20 patients who had mild, moderate or severe COVID19 were asked to donate blood 6 weeks after infection diagnosis. Patients were classified into mild, moderate and severe COVID19 based on supportive respiratory treatment. Patients with mild COVID19 did not require oxygen treatment. Patients with moderate COVID-19 required supplementary oxygen support wheras patients with severe COVID-19 required non-invasive ventilation or high-flow nasal cannula oxygen therapy. All participants gave written informed consent to participate in the study which was approved by the Swedish ethical review authority (2020/01747). Blood was drawn in citrated tubes and plasma was stored in the -80 °C. For B cell isolation and antibody discovery, patients convalescing after severe COVID19 infection donated blood 6 weeks after discharge from the hospital. Thirtyml of blood were drawn into citrated tubes and the B cells were directly isolated using Rosettesep B (according to the manufacturor’s instructions) and frozen at -150°C. B cells were harvested from 7 donors and were kept frozen until the sorting day when 10^7^ cells were thawed, pooled and prepared for baiting which was performed in PBS +2% FBS. Spike protein (S1+S2 ECD-His Recombinant Protein) was purchased from SinoBiologicals (cat: 40589-V08B1) and was reconstituted to 1 mg/ml in PBS. Spike protein was conjugated to Alexafluor 647 microscale labeling kit (Invitrogen). The fluorescently labelled spike protein was incubated with the pooled B cells at a concentration of 0.5 µg/ml for 30 mins on ice. The cells were then washed with PBS, blocked in 2% BSA and stained with antibodies against CD19-PE (BD-555413), CD3-BV510 (BD-564713), IgG-BV421 (BD-562581) and a live/Dead Sytox stain. The cells were stained for 30 mins on ice and were later washed and prepared for sorting. Bulk cell sorting was performed using a FACSAriaFusion sorter, where the gates were set using unstained and FMO-1 controls. 7000 spike-reactive cells were sorted into RPMI + 10% FBS and were transported immediately to the RNA-sequencing facility while on ice.

### 10X genomics sequencing and data analysis

We performed 10X Genomics single-cell sequencing on the 7000 Spike-reactive cells (Center for Translational Genomics facility, Lund University). Cellranger suite cellranger mkfastq was used for demultiplexing and cellranger vdj for generating V(D)J sequences and annotation. Once received, we collated the V(D)J regions from our antibodies of interest using the V-Loupe software (10X Genomics software platform). 96 antibodies were chosen based on their phylogenetic distribution and the light and heavy chain variable regions were cloned into an IgG1 expression vector (Twist Biosciences). The 192 antibody plasmids (light and heavy chain constructs) were transformed into chemically competent Mix’n’go E. coli (Zymo research, T3002) and minipreps were prepared from the resultant colonies. Multiple sequence alignment using the ClustalW algorithm was performed on the light chain sequences and the heavy chain sequences. Single-linkage clustering was performed using the sum of the Hamming distances between the aligned light chain and the heavy chain as the similarity metric.

### Antibody reactivity screening

For ELISA, 10 µg/ml of Spike protein diluted in PBS was immobilized onto ELISA wells overnight at 4°C. The wells were washed with PBST and 100 µl of antibody supernatants were added to each well. A negative control (normal human pooled serum) and positive control (COVID patient serum) were used at 10% dilutions (in PBS). After one hour of incubation at 37 °C, the wells were washed and HRP-conjugated protein G (Biorad 1706425) was added and kept for one hour at 37 °C. The wells were finally washed and developed with 100 µl developing reagent (20 ml Substrate buffer NaCitrate pH 4.5 + 1 ml ABTS Peroxide substrate + 0.4 ml H2O2). OD450 was recorded and plotted.

For bead-based screening, fluorescent (APC) streptavidin microsphere beads (1 µm, Bangs Laboratories, Cat: CFR004) were used as Spike carriers. Spike protein was conjugated to biotin using the EZ-Link™ Micro Sulfo-NHS-LC-Biotinylation Kit (Thermofischer; Cat: 21935). The biotinylated Spike protein was attached to the streptavidin microbeads according to the bead manufacturer’s instructions. For antibody reactivity testing, the Spike-beads were blocked with 5% BSA (in PBS) for 30 mins at 37°C. 150k beads were then centrifuged and incubated with 1 µg/ml of antibody for one hour at 37°C in 96-well plates. The beads were washed with PBS and a secondary Alexa Fluor 488 conjugated Fab -Fab antibody (Jackson laboratories) was used to develop fluorescent signal. After a 30 min incubation with the secondary antibodies, the beads were further washed and fluorescence was detected using a Beckman Coulter Cytoflex flow cytometer.

### Spike-THP-1 association assays

Spike-beads were opsonized with patient plasma or monoclonal antibodies at the specified concentrations for 30 minutes at 37°C in a 100 µl volume in 96 well plates. The beads were then centrifuged and reconstituted in 50 µl Sodium medium (5.6 mM glucose, 127 mM NaCl, 10.8 mM KCl, 2.4 mM KH2PO4, 1.6 mM MgSO4, 10 mM HEPES, 1.8 mM CaCl2; pH adjusted to 7.3 with NaOH). THP-1 cells were washed twice with PBS and reconstituted in Sodium medium. Spike beads and THP-1 cells were mixed at a ratio of 2 beads per THP-1 cell, in a final volume of 100 µl of Sodium medium. The suspension was mixed and cooled on ice for 5 minutes before incubating at 37°C in a shaking incubator for 30 minutes. The suspension was later cooled and analyzed via flow cytometry. Gating was first set on the cell population and the percentage of cells associated with beads (now fluorescent in the APC channel) was determined (Fig. 1a). After cell-spike reactivity analysis was done, the cells were centrifuged and fixed with methanol (for 10 minutes at room temperature). The cells were then washed and resuspended in PBS, awaiting further flow cytometry analysis. Gates were then changed to include all the beads in the APC-fluorescent channel (Fig. 1b, top right). For internalization analysis, Spike-beads were conjugated with pHrodo (FITC), an acid-sensitive dye that fluoresces in acidic environments. The beads were opsonized with different concentrations of antibodies and then interacted with THP-1 cells. Cells determined to be fluorescent in the APC and FITC channels by flow cytometry have had internalized as well as associated beads.

### Pseudotyped virus neutralization assays

Pseudotyped lentiviruses displaying the SARS-CoV-2 pandemic founder variant (Wu-Hu-1) packaging a firefly luciferase reporter gene were generated by the co-transfection of HEK293T cells using Lipofectamine 3000 (Invitrogen) per the manufacturer’s protocols. Media was changed 12-16 hours after transfection, and pseudotyped viruses were harvested at 48- and 72-hours post-transfection, clarified by centrifugation, and stored at -80°C until use. Pseudotyped viruses sufficient to generate 50,000 relative light units (RLUs) were incubated with serial dilutions of antibodies for 60 min at 37°C in a 96-well plate, and then 15,000 HEK293T-hACE2 cells were added to each well. For these experiments, the HEK293T-hACE2 cell culture was supplemented with penicillin/streptomycin antibiotics to avoid contamination. Plates were incubated at 37°C for 48 hours, and luminescence was then measured using Bright-Glo (Promega) per the manufacturer’s protocol, on a GM-2000 luminometer (Promega).

### Bead-based neutralization assay

HEK293T-ACE2 cells were seeded at density of 35,000 cells per well in a Poly-D-Lysine coated flat bottom 96 well plate. The outer skirt wells were kept cell free and were filled with medium. The day of the experiment, Spike-beads were distributed to fresh 96 well plates, adding 700,000 beads/ well. The beads were opsonized with 100 µl of antibody supernatants at 37°C for one hour. The beads were then resuspended by pipetting up and down and the bead/antibody mix was used to replace the medium on the HEK293T-ACE2 cells. The cells were incubated with beads for one hour at 37°C. The cells then were washed three times with PBS and fixed with 4% paraformaldehyde at room temperature for 15 minutes. The cells were finally washed and prepared for imaging. Four images from the center of the field of each well in the 96-well plate were acquired using 10X magnification. The number of beads per field was automatically determined using the Nikon Jobs software. For each experiment, the average number of beads/quadrant per all 96 wells was calculated and used as a 100% reference. We chose to normalize our data internally this was because our hypothesis was that the majority of our antibodies would not be neutralizing. Data from four experiments were pooled and presented.

### Animal experiments

Forty-two nine-week old female K18 hACE2 (B6.Cg-Tg(K18-ACE2)2Prlmn/J) mice were inoculated intranasally with 10^5^ PFU of SARS-CoV-2 (Wuhan strain, isolate SARS-CoV-2/01/human/2020/SWE, sourced from the Swedish Health Authorities). These mice are transgenic and carry the human ACE2 gene, making them permissive to SARS-CoV-2 infection (Jackson laboratories). One day after infection, the mice were split into 6 groups of 7 mice and antibodies were administered in one single dose intraperitoneally. We opted for a therapeutic model because we wanted to test the therapeutic potential of our antibodies under the most robust conditions. The body weights of the mice were recorded daily and the animals were euthanized if they lost more than 20% of their body weights or showed a severe deterioration in health status. The infection proceeded for 7 days before the animals were euthanized. Blood, tissue and bronchoalveolar lavage were harvested and stored accordingly. All the animal experiments were performed under the approval of the regional animal experimental ethics committee in Stockholm (16765-2020).

### Determination of IgG-antigen interaction kinetics

Analysis of RBD-IgG reaction kinetics was performed on a MASS-16 biosensor instrument (Bruker, Hamburg, Germany). Anti-Human IgG (Fc) (Cytiva, Uppsala, Sweden) was diluted to 25 µg/ml in 10 mM sodium acetate buffer pH 5 and immobilized on a High Capacity Amine Sensor chip (Bruker) (time of interaction: 7 min; flow rate: 10 µl/min). S-protein-specific IgG was diluted in running buffer (Dulbecco’s PBS (HyClone, South Logan, UT, USA) containing 0.01% Tween 20) and allowed to bind during a 90 s long injection (flow rate: 10 µl/min). Its capture level was set to be below 140 RU. The antigen (SARS-CoV-2 RBD (SinoBiological, Beijing, China; product number 40592-V08H) at 0.7-180 nM or Spike protein at 0.4-90 nM in running buffer) was subsequently injected (time of interaction: 2 min; flow rate: 30 µl/min). Dissociation was subsequently allowed to proceed for 5-15 min. The sensor chip was regenerated by treatment with 3 M magnesium chloride solution (Cytiva). All interactions were performed at 25° C. Apparent reaction rate kinetics was determined using a Langmuir 1:1 model using the Sierra Analyser software version 3.4.3 (Bruker).

### Competition ELISA to define relative epitope location

High binding polystyrene 96-well plates (Corning Inc., Corning, NY, USA) were coated with 2 µg/ml streptavidin (Thermo Fisher Scientific, Waltham, MA, USA) diluted in Dulbecco’s PBS (HyClone, South Logan, UT, USA) over night at +4°C. On the following day the plate was washed and subsequently incubated for 30 min with 30µl 30 nM biotinylated SARS-CoV-2 RBD (SinoBiological; product number:40592-V27H-B) diluted in Dulbecco’s PBS containing 0.05% Tween 20 and 0.5% fish gelatine (Sigma Aldrich, St. Louis, MO, USA) (assay buffer). After washing the immobilized antigen was preincubated for 40 minutes at room temperature with 30 µl assay buffer or assay buffer containing 4.8 pmol IgG. Subsequently, 10 µl of assay buffer or assay buffer containing 4.8 pmol scFv was added to each well. After 1 hour incubation at room temperature the wells were washed and bound scFv was detected by incubation for 40 minutes at room temperature with peroxidase labelled monoclonal anti-FLAG® M2 antibody (Sigma Aldrich (30 µl diluted 1/4000 in assay buffer) and development using 1-Step™ Ultra TMB-ELISA Substrate Solution (Thermo Fisher Scientific).

### Surface plasmon resonance studies to assess IgG-specificity

The ability of IgG to interfere with the binding of SARS-CoV-2 RBD to its receptor, Angiotensin-Converting Enzyme 2 (ACE2) was examined by surface plasmon resonance-based detection in real time using a MASS-16 instrument (Bruker, Hamburg, Germany). The spots on a High Capacity Amine Sensor chip (Bruker) were immobilized with streptavidin (ThermoFisher Scientific, Waltham, MA, USA) (50 µg/ml diluted in 10 mM sodium acetate buffer pH 5.0; flow rate: 10 µl/min; time of immobilization: 6 min) to a level of approximately 1000 RU. Subsequently 50 nM biotinylated ACE2 (SinoBiological, Beijing, Shina; product number: 10108-H08H-B) was immobilized onto the chip’s A spots (flow rate: 10 µl/s; time of binding: 2 min) while B spots were used as reference spots without ACE-2. 40 and 26 nM Receptor Binding Domain (RBD) was pre-incubated with 200 nM IgG diluted in Dulbecco’s PBS (HyClone, South Logan, UT, USA) containing 0.01% Tween 20. The mixtures were injected over the sensor chip for 2 min, followed by a 6 min dissociation phase (flow rate: 30 µl/min). The sensor chip was regenerated by treatment with 1 M magnesium chloride solution (Sigma Aldrich, St Louis, MO, USA).

Binding of IgG to different mutated versions of SARS-CoV-2 was examined by a surface plasmon resonance assay. A High Capacity Amine Sensor chip (Bruker) was immobilized with F(ab’) Goat Anti-Human IgG, Fcγ fragment specific (Jackson, Ely, UK) at 50 µg/ml in 10 mM sodium acetate buffer pH 5 (time of interaction: 7 min; flow rate: 10 µl/min). Antibodies were diluted in Dulbecco’s PBS (HyClone, South Logan, UT, USA) containing 0.01% Tween 20 and injected over the surface for 2 minutes at 10 µL/min. The antigens, produced in HEK293 cells, were obtained from SinoBiological (Beijing, China; product numbers: SARS-CoV-2 Spike RBD: 40592-V08H; SARS-CoV-2 Spike RBD-N501Y: 40592-V08H82; SARS-CoV-2 Spike RBD-E484K: 40592-V08H84; SARS-CoV-2 Spike RBD-K417N, E484K, N501Y: 40592-V08H85; SARS-CoV-2 Spike S1 HV69-70 deletion, Y144 deletion, N501Y, A570D, D614G, P681H: 40591-V08H12). All proteins were diluted to 50 nM in Dulbecco’s PBS containing 0.01% Tween 20 and injected over the surface (time of interaction: 2 minutes; flow rate: 30 µl/min) followed by a dissociation phase of 6 minutes. After each cycle the surface was regenerated with 10 mM glycin pH 2.2 containing 30 mM HCl.

### Crosslinking of antibodies to Spike protein

For the crosslinking of the antibodies to the Spike protein, 2 uG of each antibody was separately cross-linked to 2 uG of the Spike protein (Sino Biological Inc. 40589-V08H4 LC14SE2504, Recombinant SARS CoV-2 (1029-nCoV) Spike), as previously described (42). Briefly, the proteins were allowed to bind to each other in 50 uL of 1xPBS, pH 7.4 at 37 °C, 500 rpm, 15 min. Heavy/light disuccinimidylsuberate (DSS; DSS-H12/D12, Creative Molecules Inc.) resuspended in dimethylformamide (DMF) was added to final concentrations 250 and 500 µM and incubated for a further of 60 min at 37 °C, 800 rpm. The crosslinking reaction was quenched with a final concentration of 50 mM ammonium bicarbonate at 37 °C, 800 rpm, 15 min.

### Sample preparation for MS

The crosslinked samples mixed with 8 M urea and 100 mM ammonium bicarbonate, and the cysteine bonds were reduced with 5 mM TCEP (37 °C for 2h, 800 rpm) and alkylated with 10 mM iodoacetamide (22 °C for 30 min, in the dark). The proteins were first digested with 1 µg of sequencing grade lysyl endopeptidase (Wako Chemicals) (37 °C, 800 rpm, 2h). The samples were diluted with 100 mM ammonium bicarbonate to a final urea concentration of 1.5 M, and 1 µg sequencing grade trypsin (Promega) was added for further protein digestion (37 °C, 800 rpm, 18 h). Samples were acidified (to a final pH 3.0) with 10% formic acid, and the peptides purified with C18 reverse phase spin columns according to the manufacturer’s instructions (Macrospin columns, Harvard Apparatus). Peptides were dried in a speedvac and reconstituted in 2% acetonitrile, 0.2% formic acid prior to mass spectrometric analyses.

### Liquid chromatography tandem mass spectrometry (LC-MS/MS)

All peptide analyses were performed on Q Exactive HF-X mass spectrometer (Thermo Scientific) connected to an EASY-nLC 1200 ultra-high-performance liquid chromatography system (Thermo Scientific). Peptides were loaded onto an Acclaim PepMap 100 (75µm x 2 cm) C18 (3 µm, 100 Å) pre-column and separated on an EASY-Spray column (Thermo Scientific; ID 75µm x 50 cm, column temperature 45°C) operated at a constant pressure of 800 bar. A linear gradient from 4 to 45% of 80% acetonitrile in aqueous 0.1% formic acid was run for 65 min at a flow rate of 350 nl min-1. One full MS scan (resolution 60000 @ 200 m/z; mass range 390–1 210m/z) was followed by MS/MS scans (resolution 15000 @ 200 m/z) of the 15 most abundant ion signals. The precursor ions were isolated with 2 m/z isolation width and fragmented using HCD at a normalized collision energy of 30. Charge state screening was enabled, and precursors with an unknown charge state and a charge state of 1 were rejected. The dynamic exclusion window was set to 10 s. The automatic gain control was set to 3e6 and 1e5 for MS and MS/MS with ion accumulation times of 110 ms and 60 ms, respectively. The intensity threshold for precursor ion selection was set to 1.7e^4^.

## Author contributions

Conceptualization: WB, VWH, RW, JM, LM, MO, MR, and PN. Experimentation and data analysis: WB, SW, MS, AB, UH, OL, HK, MG, MW, EE, LH, OA, JKA, TH. Writing original draft: WB and PN. All authors contributed to reading and editing the final manuscript.

## Acknowledgements

WB, LM, JM and PN are funded by the Knut and Alice Wallenberg Foundation. TH and equipment were funded by IngaBritt och Arne Lundbergs Forskningsstiftelse. WB, MO, JM, MR, and PN were funded by grants from the SciLifeLab National COVID-19 Research Program, financed by the Knut and Alice Wallenberg Foundation. WB were funded by the Royal Physiographic Society. HK were funded by Swiss National Science Foundation (grant no. P2ZHP3_191289). We thank Åsa Petersson for help with flow sorting and Berit Olofsson for technical assistance. We thank Benjamin Murrell and Daniel Sheward at the Karolinska Institutet for help with the pseudovirus neutralization assay. We thank Ali Mirazimi at the Public Health Agency of Sweden and Karolinska Institutet, for providing virus for the animal infection experiments performed. We thank the Lund University Bioimaging Centre (LBIC) for use of fluorescence microscopes.

